# Compositional shifts associated with major evolutionary transitions in plants

**DOI:** 10.1101/2022.06.13.495913

**Authors:** Stephen A. Smith, Nathanael Walker-Hale, C. Tomomi Parins Fukuchi

## Abstract

Heterogeneity in gene trees, morphological characters, and composition has been associated with several major clades across the plant tree of life. Here, we examine heterogeneity in composition across a large transcriptomic dataset of plants in order to better understand whether locations of shifts in composition are shared across gene regions and whether directions of shifts within clades are shared across gene regions.
We estimate mixed models of composition for both DNA and amino acids across a recent large scale transcriptomic dataset for plants.
We find shifts in composition across both DNA and amino acid datasets, with more shifts detected in DNA. We find that Chlorophytes and lineages within experience the most shifts. However, many shifts occur at the origins of land, vascular, and seed plants. While genes in these clades do not typically share the same composition, they tend to shift in the same direction. We discuss potential causes of these patterns.
Compositional heterogeneity has been highlighted as a potential problem for phylogenetic analysis, but the variation presented here highlights the need to further investigate these patterns for the signal of biological processes.

**Plain language summary:** We demonstrate that many nucleotide and amino acid compositional shifts in plants occur at the origins of major clades and while individual genes do not share the same composition they often shift in the same direction. We suggest that these patterns warrant further exploration as the signal of important biological processes during the evolution of plants.

## Introduction

Heterogeneity in the patterns and processes of molecular evolution is common through time and between lineages. For example, topological conflict between different gene regions has been demonstrated to be common across the tree of life, reflecting, in part, population processes including introgression and incomplete lineage sorting (Maddison, 1997; Rokas et al., 2003; Smith et al., 2015). High rates of morphological change has also been associated with conflict at several major clades across the plant tree of life (Parins-Fukuchi et al. 2021; Stull et al. 2021). An additional widely recognized form of heterogeneity is in composition: changes in the proportion of different states, such as nucleotide bases or Amino Acids (AAs), between lineages and through time, which emerges from the interplay between mutation, gene conversion, drift and selection (Eyre-Walker & Hurst, 2001; Lynch, 2007). Compositional differences are also expressed at the site-level with different protein sites preferring different AAs (Lartillot & Philippe, 2004; Wang et al., 2008; Le et al., 2008), and genome-wide with different composition between different regions within the same genome (Lynch, 2007). Different lineages are also known to favor different synonymous codons, leading to compositional bias at the codon level (Chen et al., 2004; Plotkin & Kudla, 2011). These differences are tree-heterogeneous and interactive, so that different sites and loci might experience different compositions in different lineages at different times.

Research intersecting composition and phylogenetics has typically focused on the impact of heterogeneous composition on error in phylogenetic inference, identifying how clade-specific biases in nucleotide base composition can produce false groupings of evolutionarily distant but compositionally similar taxa (Foster, 2004; Cox et al. 2014; Cox, 2018; Sousa et al., 2020). Another less well-explored avenue is the ability for heterogeneity in composition to provide a window into the molecular and population processes impacting the genome. A separate body of research has addressed the role and influence of these processes on genomes in multiple clades (Duret & Galtier, 2009; Glemin et al., 2014; Weber et al., 2014; Clément et al., 2015; Clément et al., 2017). Mutation pressure is thought to explain some genomic patterns (Lynch, 2007), such that changes in composition might reflect important shifts between the balance of mutation and drift, and hence effective population size. GC-Biased Gene Conversion (gBGC), where GC alleles act as the donor more often than expected during recombination-associated gene conversion events, also influences genome-wide GC content. Furthermore, due to gBGC, changes in recombination rate might therefore change compositions across the tree (Marais et al., 2004; Duret & Galtier, 2009; Muyle et al., 2011; Weber et al., 2014). Changes in effective population size might drive changes in composition via an increase in the efficacy of gBGC (Weber et al., 2014). Because gBGC occurs during meiosis, increases or decreases in generation time could change composition both by changing mutation rate and changing the number of meiotic, and hence the number of gBGC, events (Romiguier et al., 2010; Weber et al., 2014).

While demographic processes may influence molecular composition, several non-demographic processes also potentially contribute to compositional change (Clément et al., 2017; Hershberg & Petrov, 2008). Selection on codon usage for translational accuracy and efficiency could explain compositional changes (Hershberg & Petrov, 2008; Qiu et al., 2011). Compositional bias itself may impact codon usage and eventually AA preference (Foster et al. 1997, Singer and Hickey 2000, Knight et al., 2001; Qiu et al., 2011). Bias in the selection for particular AAs can influence composition (Błażej et al., 2017). Compositionally mediated changes in codon usage might also influence gene expression (Zhou et al., 2016). In addition to these microgenomic processes, macrogenomic changes, such as Whole-Genome Duplication (WGD) and biased retention or loss, could also create dramatic changes in composition (McGrath et al., 2014; Veleba et al., 2014).

In plants, empirical patterns in various clades, such as the GC-richness of Commelinid monocots, have been described and explained by mutation, selection, and gBGC (Qiu et al., 2011; Serres-Giardi et al., 2012; Glemin et al., 2014; Clément et al., 2015; Clément et al., 2017). Because shifts in base composition bias can be linked with such crucial evolutionary parameters as generation time and population size, they may also shed light on major evolutionary transitions in the plant tree of life.

Models of molecular evolution typically consist of two components: relative transition rates between states, and the composition of those states. State compositions of nucleotides or AAs are typically modeled at equilibrium, assuming a process that does not vary between sites or across time (Yang, 2014). These assumptions can be relaxed in several ways including partitioned models (Lanfear et al., 2012), models that allow the equilibrium composition to vary across sites (Lartillot & Philippe, 2004; Le et al., 2008), models that vary across the tree (Galtier & Gouy, 1998; Foster, 2004), or methods that vary substitution models and compositions across branches (Jayaswal et al., 2011; Zou et al., 2012; Jayaswal et al., 2014). Phylogenetic inference can be sensitive to composition biases across clades, with conflicting resolutions drawn from homogeneous vs heterogeneous models. As a result, methods relaxing these assumptions have been a major focus for phylogenetic inference of ancient nodes across the tree of life (Sousa et al., 2020; Redmond & McLysaght, 2021; Li et al., 2021). However, if molecular and population processes are driving the patterns accounted for by heterogeneous phylogenetic models, these models could be used to detect the signal of changing evolutionary processes across the tree.

Instead of focusing on the resolution of relationships within plants, we concentrate on examining the extent to which there are compositional shifts across nodes and gene regions. One shortcoming to the application of phylogenetic methods to the detection of compositional shifts is that tree-heterogeneous methods typically require the branches of interest to be specified a priori. Consequently, several efforts have been made to relax this restriction, such as testing all branches in the tree, or by investigating summary statistics of the substitution process, or other methods (Blanquart & Lartillot, 2006, 2008; Dutheil et al., 2012). Alternatively, Bayesian MCMC jump methods have been developed that allow for uncertainty in the number and placement of shifts in composition (Foster, 2004; Gowri-Shankar & Rattray, 2007). However, computational methods that allow for integrating over the uncertainty of their placement are too burdensome for large genomic datasets with hundreds of taxa and hundreds of gene regions. In parallel, research has focused on detecting shifts in the rate of diversification or phenotypic evolution across the tree (Alfaro et al., 2009; Uyeda & Harmon, 2014; Mitov et al., 2019). One such class of method uses stepwise model selection with information criteria to automatically partition the tree into different regimes (Alfaro et al., 2009; Mitov et al., 2019), but such approaches are not commonly applied to molecular data (but see Dutheil et al., 2012).

Here, we extend methods that allow composition to vary across the tree by implementing an algorithm that detects compositional shifts by comparing models of different dimensions using information criteria. We apply our method to a large collection of orthologs of coding regions from across the Viridiplantae clade (Leebens-Mack *et al*., 2019) and, instead of targeting the impacts of composition on topological resolution, we focus on identifying compositional shifts on individual gene regions.

## Methods

### Dataset

We analyzed the nucleotide and AA data from the 1KP transcriptome project data release available at https://github.com/smirarab/1kp (Leebens-Mack *et al*., 2019) to identify patterns in compositional heterogeneity across plants. For nucleotide data, we used the “unmasked and FNA2AA” data and filtered for columns containing at least 10% of data using pxclsq from phyx (-p 0.1, Brown *et al*., 2017). We chose these alignments instead of those for which trees were already inferred in order to include third codon positions for composition analyses. We ran an analysis to detect compositional shifts in both the nucleotide (the cleaned alignments of all three codon positions and our inferred trees) and AA data (using the available alignments and trees). For these alignments, we conducted phylogenetic analyses using IQ-TREE v1.6.6 (Nguyen *et al*., 2015) under the GTR+G model of evolution. For AAs, we used the “masked FAA’’ data and the corresponding trees inferred as part of the original study. We analyzed the AA using the JTT model of evolution. We used a GTR+G model and so there could be phylogenetic error introduced from violations of homogeneous composition bias. While this may impact some edges, we have also demonstrated that our method for identifying model shifts is robust to this (see Supp. Fig. 2).

Because of the non-homogeneity of the compositional model, our analysis required rooted trees. Perfect rooting was not required and would have been prohibitive considering the variation and non-monophyly of many taxonomic groups in each gene tree (see Supp. Fig. 1). In order to accommodate this, we rooted using pxrr from phyx, applying the ranked option (-r) with the following taxa in order (taxon codes from https://github.com/smirarab/1kp/blob/master/misc/annotations.csv): UNBZ, TZJQ, JGGD, HFIK, YRMA, FOMH, RWXW, FIKG, VYER, LDRY, VRGZ, ULXR, ASZK, JCXF, QLMZ, FSQE, DBYD, VKVG, BOGT, JQFK, EBWI, FIDQ, QDTV, OGZM, SRSQ, RAPY, LLEN, RFAD, NMAK, VJED, LXRN, APTP, BAJW, IAYV, IRZA, MJMQ, ROZZ, BAKF. The ranked option searches through the list of taxa and roots on the first one present.

### Detection of compositional heterogeneity

We developed an algorithm to detect locations of shifts in stationary frequencies in state composition that we describe below (see Figure 1). The method is generalized to any state model, and so proceeds in the same way for nucleotides or AAs. It requires a rooted tree and matching alignment as input. First, the method estimates a maximum likelihood root composition for the entire dataset. Next, the tree is traversed in a postorder fashion (from the tips to the root), and a maximum likelihood composition is estimated for the subtree subtending each node, if that subtree contains more than a user-specified minimum number of tips. In this work, we considered any subtree containing at least 10 tips. Using this composition for the focal node and subtree, and the root composition for the remainder of the tree, we calculate a likelihood and the Bayesian Information Criterion (BIC: Schwarz, 1978). Once a model for every eligible subtree has been estimated, we order subtrees by their BIC (i.e., by their relative improvement in fit over the base model), add them to the model configuration, calculate a new likelihood and BIC for the whole tree and add the sub-model if the new BIC is lower (i.e., the model provides a better fit). To improve computational efficiency, we discard models if their BIC score is greater than the current model by an arbitrary cutoff (we assigned a cutoff of 35). Our method has been implemented in both Golang (for flexibility) and C (for speed), and the source code is available at https://git.sr.ht/~hms/janus and https://git.sr.ht/~hms/hringhorni, respectively. A diagram is presented in Figure 1 and an empirical example is presented in Supp. Fig. 3.

**Figure 1.**
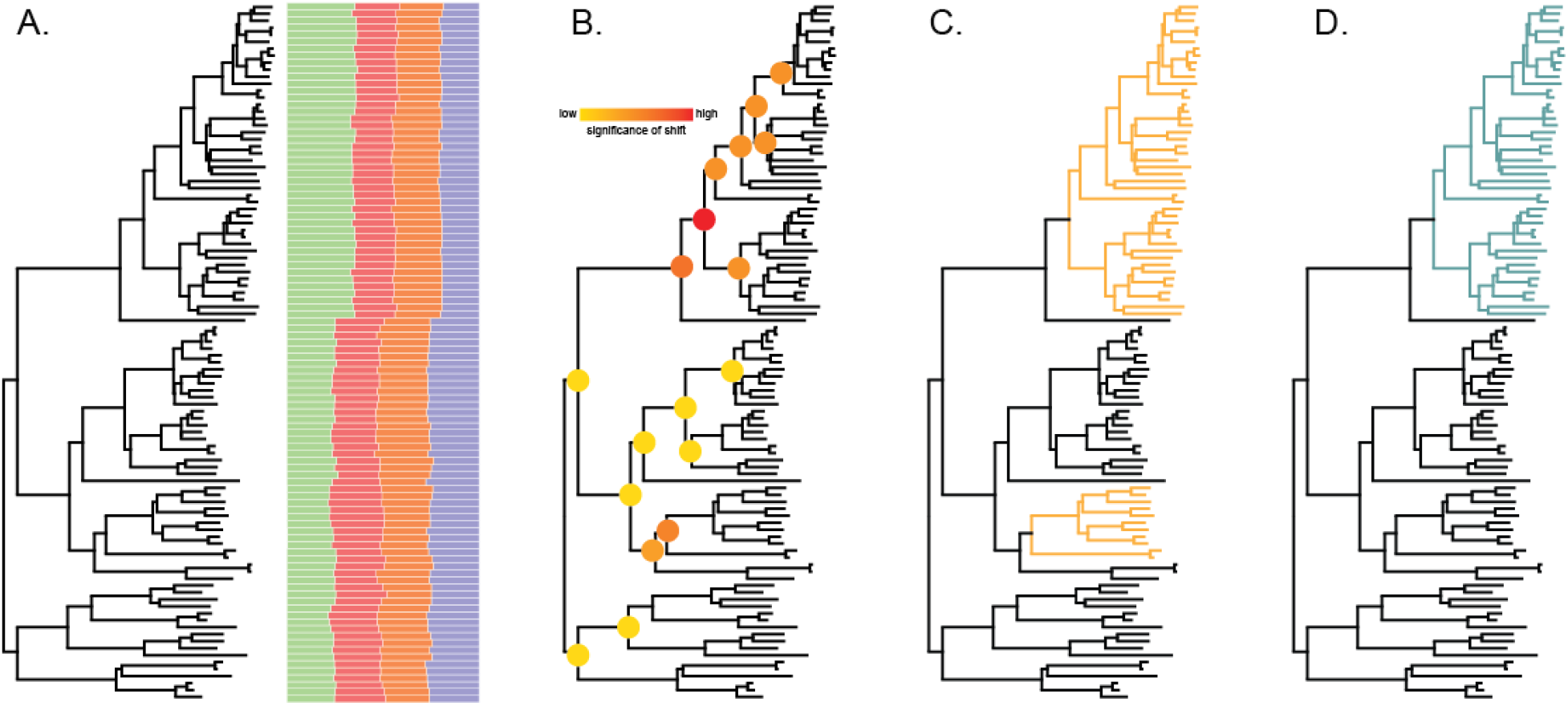
A demonstration of the procedure introduced here used on each gene tree. A) shows a tree and the sequences to the right represented as their composition of DNA. B) is the same tree with node colors corresponding to the IC values sorted with red being the highest and yellow being the lowest. C) identifies two clades as having potential shifts with only one supported after uncertainty analyses (the blue clade in D).

### Accommodating model uncertainty

One common challenge in information criterion (IC) based approaches to model comparison is their tendency to overfit, sometimes favoring models of higher complexity than the generating model. Our solution to this tendency was to assess statistical uncertainty in each model shift by estimating the relative support for the model that includes the shift vs the model without the shift. We performed these tests using BIC weights *(wBIC)*, comparing, for each putative shift, the BIC of the full model containing all inferred shifts to one dropping each individual model shift. The strength of support for each inferred shift was thus calculated by calculating the relative BIC of each candidate model *i* (in this case, shift vs no shift):

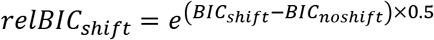

And assessing support for the shift as the ratio of the ratio of that model over the sum of all *i* candidate models:

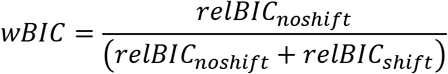

This calculation yields an index between 0 and 1, where values closer to 0 indicate weaker support for the shift, and values closer to 1 indicate stronger support. Using the reasoning that spurious shifts will likely typically be poorly supported, we removed shifts with wBIC support values below 0.95.

### Simulations

We conducted several simulations to validate the performance of our algorithm in detecting model heterogeneity. Phylogenies were simulated under a birth-death model with phyx using the pxbdsim command with defaults, except varying the size of the tree between 100 and 250 tips, and root height set to 0.75 with pxtscale (-r 0.75) from phyx. Nucleotide and AA alignments were simulated using a simulator STONE (https://git.sr.ht/~hms/stone) that allows for shifts in composition across the tree. For nucleotides, we conducted two simulations: one under JC+G and another GTR+G (both with α = 1 for rate heterogeneity). For AAs we conducted one simulation under JTT with no rate variation. Each of these simulations had a single randomly positioned compositional shift per tree. Phylogenies were then reconstructed with IQ-TREE under the GTR+G model of evolution for nucleotide alignments and the JTT+G model for AA alignments. For each simulation set, we simulated 100 replicates. Alignment lengths were 1000 for nucleotides and 300 and 1000 for Aas.

### Summarizing compositional heterogeneity

We summarized the results from the empirical analyses in several ways. Directly comparing model shifts across genes was complicated by extensive gene tree conflict. We compared the distribution of model shifts by pairwise comparison of tips on the species tree inferred in the original paper (Leebens-Mack *et al*., 2019), recording the number of times that two tips were descended from a node with a shared model, and plotted this in a heatmap on the species tree (Supp. Fig 4). Secondly, we defined major clades in the species tree, and recorded to which groups each tip descending a model shift in each gene tree belonged. We counted the number of tips from each taxonomic group, and further counted the number of tips within those taxonomic groups which were not included in the model shift (i.e., either the model shift occurred nested within that group, or those tips were placed polyphyletically in the tree due to conflict). We manually assessed these mismatches and the position of the model shift on the gene tree and assigned the shift on the species tree to occur either i) at the node defining a major clade (assuming mismatching tips are errors), which we summarize as occurring at the origin of the clade or ii) descending a node defining a major clade, which we summarize as occurring within the clade. For individual genes, we plotted model shifts on the tree and changes in parameter estimates between models. To characterize the direction and size of parameter shifts, we used a Principal Components Analysis where each row was a single sequence and each column was the frequency of one state for that sequence (i.e., 4 columns for nucleotides and 20 for Aas). We projected every gene tree onto the same set of axes for the first two PCs and colored each point (representing a single tip), by the model from which it was descended. We characterised shift direction and size by projecting fitted model parameters onto the same PC space, and calculating the vector direction and magnitude between the two sets of coordinates representing the parent and descendant model.

## Results

### Simulations

Our simulations demonstrate that, given sufficient data (i.e., alignments of sufficient length), our method has acceptable false positive and negative rates (Table 1). False positive rates were negligible after removing shifts that were poorly supported by BIC. In general, we consider the false positive rates to be of more concern than false negatives rates, but the latter were also negligible in our simulations. The highest rates of false positives were observed in short (300 site) AA alignments, which were diminished but not entirely alleviated by taking uncertainty into account. False positive rates were generally elevated when tree reconstruction error existed in the simulated data. Our simulations also demonstrate that phylogenetic reconstruction error, as measured by average RF between the simulated and reconstructed trees, occurred under each condition, including with 0 shifts. The RF distance of phylogenies that have one shift with 100 tips and zero shifts with 100 shifts are not significantly different. Therefore, instead of corresponding to the number of shifts or the presence of compositional bias, these errors seem to correspond to tree size. We also demonstrate that shifts can be identified correctly even when the phylogeny was reconstructed incorrectly (see Supp. Fig 2).

**Table 1.** Results of simulations for both nucleotide (JC/GTR) and amino acid data. Shown are false positive (False +) with and without considering uncertainty (unc). We also show results considering the correct tree and the tree based on reconstructions (rec). Finally, we present the average RF distance between the reconstructed trees and the true tree.

| # sh | # tips | N/A | Len | False + | False +<br>unc | False<br>+(rec) | False<br>+(rec)<br>unc | False - | False<br>- unc | False -<br>(rec) | False<br>- (rec)<br>unc | Avg. RF |
| --- | --- | --- | --- | --- | --- | --- | --- | --- | --- | --- | --- | --- |
| 0 | 100 | N | 1000 | 0/0.02 | 0/0 | 0/0.01 | 0/0 | - | - | - | - | 9.96/10.88 |
| 1 | 100 | N | 1000 | 0/0.04 | 0/0 | 0/0.04 | 0/0.01 | 0/0 | 0/0 | 0/0 | 0/0 | 8.76/10.16 |
| 2 | 150 | N | 1000 | 0.14/0.13 | 0.03/0.01 | 0.09/0.14 | 0/0.04 | 0/0.04 | 0.02/0.04 | 0/0.05 | 0.02/0.05 | 15.0/16.84 |
| 2 | 250 | N | 1000 | 0.1/0.14 | 0.01/0.01 | 0.1/0.12 | 0.02/0.03 | 0.01/0.04 | 0.03/0.05 | 0.04/0.06 | 0.07/0.08 | 24.8/26.34 |
| 0 | 100 | A | 300 | 0 | 0 | 0 | 0 | 0 | 0 | 0 | 0 | 14.32 |
| 1 | 100 | A | 300 | 0.02 | 0.01 | 0.11 | 0.07 | 0 | 0 | 0.02 | 0.02 | 15.9 |
| 2 | 150 | A | 300 | 0.03 | 0 | 0.18 | 0.07 | 0.01 | 0.01 | 0.01 | 0.01 | 21.34 |
| 2 | 250 | A | 300 | 0.02 | 0 | 0.19 | 0.10 | 0.02 | 0.03 | 0.03 | 0.01 | 35.6 |
| 0 | 100 | A | 1000 | 0 | 0 | 0 | 0 | 0 | 0 | 0 | 0 | 4.84 |
| 1 | 100 | A | 1000 | 0.01 | 0 | 0.03 | 0.01 | 0 | 0 | 0 | 0 | 4.76 |
| 2 | 150 | A | 1000 | 0.18 | 0 | 0.19 | 0 | 0 | 0 | 0.01 | 0.01 | 6.82 |
| 2 | 250 | A | 1000 | 0.22 | 0 | 0.22 | 0.01 | 0 | 0 | 0 | 0 | 12.0 |

### Phylogenetic patterns of compositional shifts

We applied our method to a large dataset of orthologs derived from genomes and transcriptomes across Archaeplastida. As noted in the original study (Leebens-Mack et al., 2019), the inferred gene trees contained high levels of conflict. For example, 38% of nucleotide and 32% of AA gene trees contained non-monophyletic seed plants. We searched for compositional shifts in inferred gene trees from nucleotide and AA data. We detected multiple shifts in both datasets, with many more shifts detected for nucleotide data (**Figure 2**). The phylogenetic location of these shifts differed between different trees, and we observed a great deal of gene tree conflict between the individual orthologs and the species tree, complicating the localization of shifts. Nevertheless, general patterns did emerge when comparing shift locations to the species tree (**Figure 2**). Many nucleotide shifts were detected at the Embryophyta node, corresponding to the origin of land plants, at the Tracheophyta node corresponding to the evolution of vascularity, at the node uniting ferns and the rest of Spermatophyta, at ferns, at the Spermatophyta node corresponding to the evolution of seeds, and at the Angiosperm node corresponding to the evolution of flowers. Many nucleotide shifts were also detected at the base of and within Chlorophytes. By contrast, AA shifts were enriched at the Spermatophyta and Angiosperm nodes and were similarly common at and within Chlorophytes. Several shifts were identified within the named clades, such as at or within Eudicots, could not be explored further because our sampling or the conflict in the gene tree precluded further localization.

**Figure 2.**
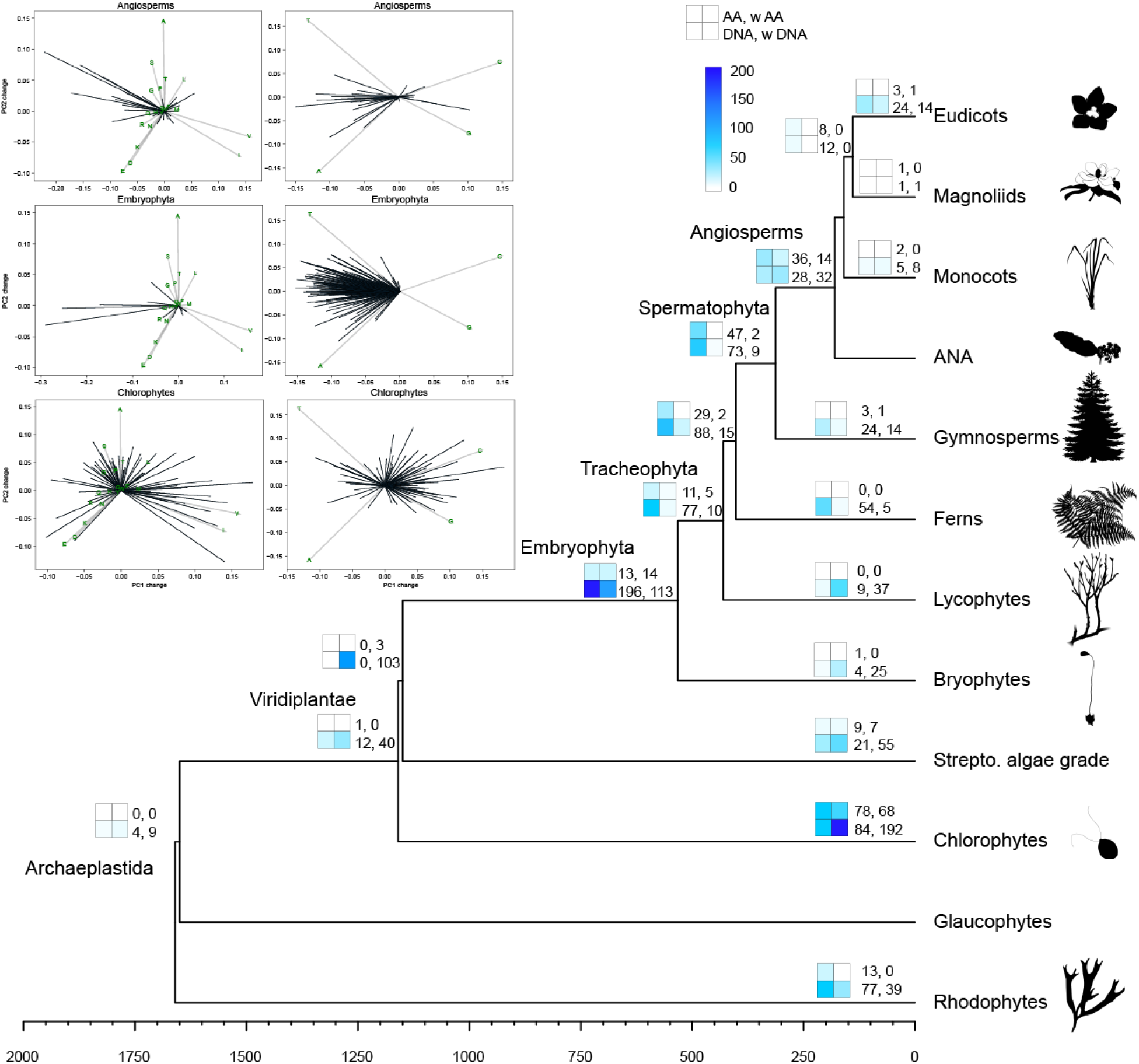
Summarized results for AA and DNA. Inset plots denote vectors of composition shifts for both AA (left) and DNA (right) for Angiosperms, Embryophyta, and Chlorophytes. For the complete set, see Supp Figs. 5 and 6. The black lines in each plot represents a single shift within a single gene. The direction shows the composition shift (e.g., most of the shifts in Embryophyta DNA plots shift to more A and T) and the length of the line shows the strength of the shift. The phylogeny on the right shows shifts detected by clade. There are four boxes at each major clade that correspond to, starting from top left to bottom right, shifts in AA data at that node, shifts in AA data within that node (e.g., because the clade was not monophyletic or because the shift is missing one or more taxa within the clade), shifts in DNA data at that node, and shifts in DNA data within that node. Colors correspond to the number of shifts. For example, at Embryophyta, there are 196 DNA shifts at that node and 113 shifts that occur within that node (missing one or more Embryophyta but not so many as to be considered Tracheophyta or Bryophytes).

**Figure 3.**
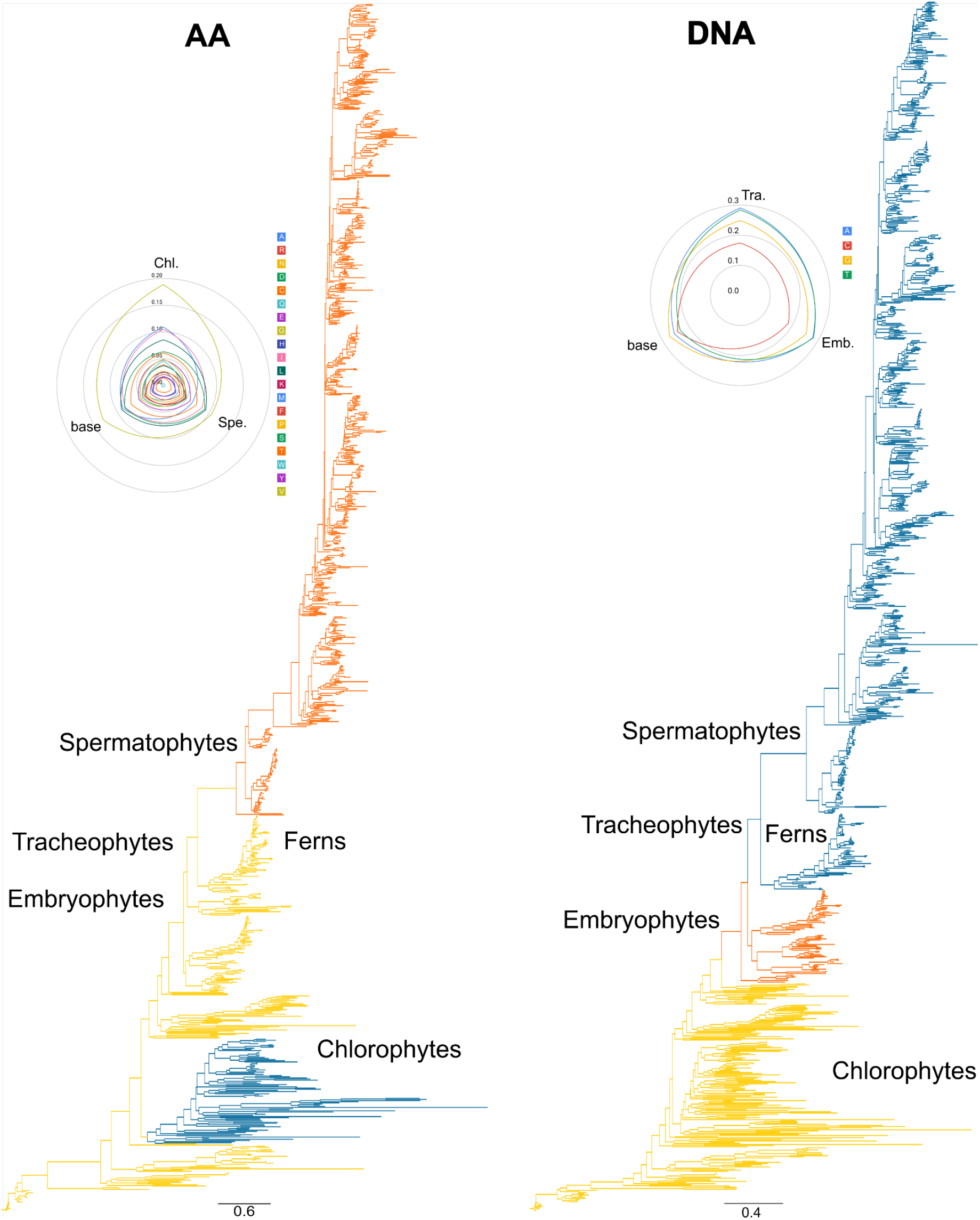
Ortholog 5936 results from both AA and DNA datasets. Colors are meant to identify shifts within the dataset (shared colors between AA and DNA datasets do not denote shared models between AA and DNA results). Base composition model results are presented in radar graphs where lines represent the proportion of the composition in each amino acid or base. For example, in comparing Tracheophytes and Embryophytes to the base model for DNA, there is an increase in As and Ts.

### Direction of compositional shifts

The direction of compositional shifts (i.e., which state frequencies increased or decreased between a parent and child model) differed both within and between genes. While specific compositional values may not be shared by many genes, we noticed a tendency for shifts at comparable nodes to occur in similar directions (**Figure 4**). The root nodes of angiosperms, chlorophytes, and embryophytes each displayed many nucleotide composition shifts that were, for angiosperms and embryophytes, heavily directionally biased towards higher AT (**Figure 2**). Several nodes displayed similarly biased amino acid compositional shifts. These biased shifts were highly evident at the origin of Tracheophyta, angiosperms, Zygnematophyceae, Spermatophyta, Embryophyta, and chlorophytes (Supp. Figs. 5-6).

**Figure 4.**
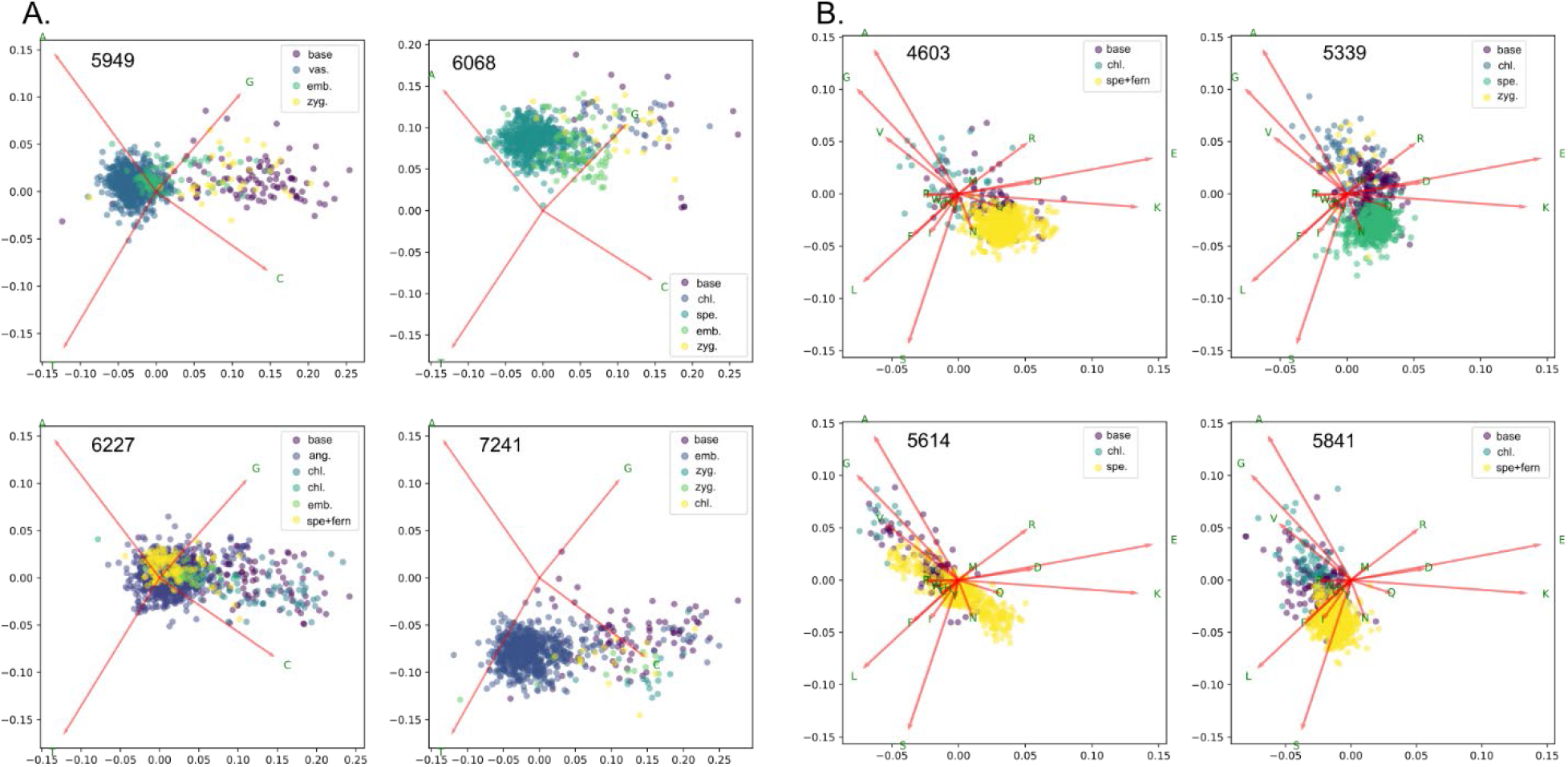
Principal component analyses of four DNA datasets (A) and four AA datasets (B) with each point representing one taxon and colors denote shared shifts within the dataset. PC loadings are based on the entire DNA and AA datasets respectively to allow for easier interpretation. For 5949, vascular plants and embryophytes have more AT bias than tips sharing the base model. The same pattern is seen for 6068 for spermatophytes and embryophytes, angiosperms and spermatophytes in 6227, and embryophytes in 7241. While each is shifting to more AT, given that these are plotted with the same PC loadings, they are also not converging on the same space.

To determine whether patterns in the direction of nucleotide compositional shifts were related to codon usage bias, we examined codon usage for each model within each gene. We noted several patterns. Firstly, codon usage was strongly biased within each residue, and there is a tendency for land plants to feature more AT-rich codons. Additionally, clades nested within land plants (e.g., Embryophyta, Tracheophyta) tend to be more AT-rich than other clades (e.g., Bryophytes). Gymnosperms showed the highest degree of codon usage bias, favoring AT-rich codons.

## Discussion

The results of the analyses of the direction of the compositional shifts and the phylogenetic position of the shifts suggest a common or related causes for these biases for major clades of land plants. The most notable pattern in this dataset is the tendency for compositional shifts of Embryophytes, Tracheophytes, and Spermatophytes to be shift to be more AT enriched. Many of these compositional shifts occur at the origins of these major named clades. The primary goals of this study are to demonstrate notable patterns of compositional shifts across vascular plants across gene trees, where previously research has focused on the accuracy of phylogenetic reconstructions using heterogeneous composition. We discuss potential causes of this heterogeneity and where certain causes seem plausible based on the analyses here as well as previous studies. However, additional lines of evidence will be necessary to further narrow these causes. Nevertheless, the patterns presented here are substantial enough to warrant further investigation.

### Life history

In our analyses, Chlorophytes tend to have shifts in compositional vectors that vary widely, some shifts toward elevated GC and some toward elevated AT (Figure 2). In contrast, land plants, vascular plants, seed plants, and flowering plants, tend to show, when there are shifts in composition, a tendency towards stronger AT bias. Furthermore, while these genes show trends towards more AT, there is not a clear lineage specific optimal AT. In other words, each gene increases in AT but not to the same AT across genes, which reflects documented intragenomic variation in base compositions (Clement et al., 2017; Glemin et al., 2014). There may be many potential causes for these patterns, however, one notable difference between those lineages with shifting AT bias are dramatic changes to life history. Life history has been demonstrated to have an impact on genome composition. For example, biased gene conversion can favor the proliferation of GC alleles during meiotic recombination, such that short generation time could lead to increased GC-richness (Duret & Galtier, 2009; Weber et al., 2014). On the other hand, mutation tends to be AT biased and lineages with longer generation times are expected to have higher mutation rates due to more cell divisions and accumulated DNA damage (Lynch, 2007, Bergeron et al. 2023). Population size also plays a compounding role. Large effective population sizes tend to make natural selection more effective, and in the case of composition bias this may translate into composition reflecting advantageous selection more than bias. On the other hand, smaller effective population sizes increase the probability that mutations will be fixed by drift. Large population sizes and increased generation times are associated with higher equilibrium GC and faster increases of GC content (Romiguier et al., 2010), suggesting that reductions in equilibrium GC might reflect shrinking effective population sizes or increased generation times. Our demographic model suggests that changes at land plants, vascular plants, seed plants, and angiosperms moved lineages closer to mutation-drift equilibrium and away from strong natural selection and BGC (Clement et al. 2017). For Chlorophytes with short generation times and larger population sizes, this may reflect the variable gene composition. Of note, are the gymnosperms which tend to have higher composition bias but fewer phylogenetic shifts. Our failure to detect shifts, however, may be due to lower taxon sampling of the gymnosperms. Alternatively, the slower generation time of gymnosperms may also play a role, which may have prevented them from reaching compositional consistency between lineages (Lanfear et al., 2013). This would yield weaker signals for our methods to detect shifts.

Our expectations under a model of mutation bias is that populations with slower generation time and smaller effective population sizes will have lower GC-richness and higher AT-richness at equilibrium because of AT-biased mutations and a lower rate and a lower efficiency of gBGC. Our results are consistent with many major changes in traits and life history across the Viridiplantae being associated with longer generation times and/or reductions in effective population size. This pattern seems likely to be true of gymnosperms, which are large, long-lived trees with slow generation times (De La Torre et al. 2017) and our results suggest that it is true of angiosperms and other lineages.

### Selection

In contrast to the demographic explanation above, selection might also drive the evolution of base composition (Clement et al., 2017; Qiu et al., 2011). Selection on codon usage could lead to preferred codons for given amino acids which are more GC- or AT-rich, leading to genome-wide patterns (Hershberg & Petrov, 2008). Because of the bias in codon composition for certain amino acids, shifts in amino acid preference at particular sites could also produce a compositional impact (Jobson & Qiu, 2011, but see Wang et al., 2004). In an analysis of extant plant genomes, Clement et al. (2017) found that the role of selection on codon usage in driving composition was small relative to BGC. However, we cannot rule out that selection played a role in generating the patterns we observe here. Moreover, these two explanations are not mutually exclusive. Selection is expected to be more efficacious in larger populations, so the possible demographic changes we suggest might interact with selection to produce changes in equilibrium composition. Further population genetic analysis of extant populations will be necessary to inform the degree to which these processes interact to shape natural variation in base composition, including in response to changing population size, generation times, or major modes of life history (Qiu et al., 2011b). Due to the necessarily coarse nature of our investigation, it is difficult to comment on how different processes might contribute to the patterns we observe. Such a distinction is a goal of further modeling efforts (Kostka et al., 2012), and will undoubtedly be important in more focused studies of single organisms or loci.

### Population processes, base composition, and gene tree discordance

Base compositional biases have been hypothesized to be linked to numerous explicit population processes, including those outlined above. We suggest that the patterns in base composition shifts that occur at key nodes in plant phylogeny are likely the result of some combination or subset of these, and perhaps other, population processes. For example, while we expect life history shifts, such as lengthening of generation time, to correspond to increases in AT-content, it is important to note that this pattern may also be consistent with myriad other lower-level processes. Empirically demonstrating a robust link between such broad-scale patterns as those explored here to specific population processes is notoriously challenging in macroevolutionary studies. In this study, we were focused on harnessing our new approach on pattern discovery first, while also considering some possible explanations for these patterns at the population level. Future work will be needed to more explicitly distinguish between these candidate processes and understand how each maps to broadly-observable phylogenetic patterns, such as those reconstructed here. For now, we lack a rigorous understanding of how specific population processes scale up to phylogenetic patterns and so the first step is to consider as many candidate processes as possible. A first step may be to identify whether life history shifts are *statistically* linked with differential patterns in AT-richness. Moving forward, it will become important to better understand how and whether population processes can be statistically identified from one another from phylogenetic patterns. Nevertheless, the timing of base composition shifts that we identify here suggests that major plant clades are reflective of fundamental biological revolutions, with effects spanning organismal scales from the genome, through life history, and morphology (Donoghue 2005).

One increasingly common avenue through which to explore population dynamics such as incomplete lineage sorting (ILS) and introgression is to explore patterns in gene-tree conflict (Smith et al. 2015; Smith et al. 2020). We observed substantial topological discordance between the gene trees analyzed. It has been previously suggested that biases in base composition may drive error in species tree reconstruction (Cox 2018, Foster 2004). In principle, it is possible that some proportion of the extensive topological conflict we found in the present dataset was caused by differential base composition bias across the loci. However, Robinson-Foulds distances between each gene tree and the species tree were primarily correlated with tree size with a weak correlation to the number of inferred composition shifts in nucleotides, but a weak negative relationship for AAs, and a great deal of variance unexplained (Table 1 and Supp Figs. 6-7). Here, at most of the major nodes we explored, we found base composition evolution to be highly biased in its direction, with most loci shifting in a similar direction. As a result, any reconstruction error caused by base composition issues would likely affect reconstruction at these nodes roughly uniformly. While we tended to observe a distribution of alternative tree topologies at each node, previous analyses have found that some of these patterns follow expectations under population processes such as ILS and introgression (Smith et al. 2020). This suggests that gene-tree discordance in this dataset is likely caused by a combination of population processes, such as ILS, and systematic error, perhaps including erroneous ortholog identification, assembly, and/or contamination. Additionally, we would expect that compositionally-driven discordance would manifest by uniting clades with disparate compositions, which our method would then tend to infer as a single, unidirectional shift, as opposed to the multiple separate shifts we observe here. Therefore, if compositionally-driven discordance is a major factor in our dataset, it should tend to make our findings conservative by reconstructing fewer shifts.

### Phylogenetic resolution

The simulations conducted here demonstrated that our method can correctly identify the location of phylogenetic shifts even in the face of reconstruction error. Nevertheless, the impact of compositional bias on phylogenetic reconstruction has been well demonstrated. The phylogenetic resolution of several deep nodes differs between genes in the DNA and amino acid datasets, and some shifts associated with deep nodes are associated with those alternative resolutions of major clades. For example, in many genes, the Bryophytes are non-monophyletic and shifts are associated with the nodes surrounding this conflicting relationship. This has been found previously by Cox et al. (2014). In gene region 6401, the Bryophytes form a grade with a shift shared by a clade of liverworts and the rest of vascular plants. The amino acid phylogeny of the same gene has no significant shift in the molecular composition. Other examples include lycopods sister to ferns versus ferns sister to seed plants– the latter is associated with shifts in molecular evolution 29 times in amino acids and 68 times in nucleotides. While the analyses presented here are not focused on the phylogenetic resolution of these major clades, other studies have demonstrated that heterogeneity can alter phylogenetic reconstruction (CITATIONS). The analyses here underscore the importance of that consideration in future studies.

### Data quality

The datasets we used here present several challenges that may stem from quality-control issues that are common among large and complex genomic datasets. We note this problem primarily because as many new genomic and transcriptomic datasets become available, as in this study, researchers will be tempted to address large scale questions taking advantage of these enormous datasets. However, caution should continue to be exercised, because errors in homology or contamination are likely still prevalent, despite researchers’ best efforts. For example, 38% of the nucleotide gene trees and 32% of amino acid gene trees have non-monophyletic seed plants. This presents several challenges, but primarily, in summarizing the phylogenetic placement results, we had to accept that there may be outlying taxa that make strict monophyly difficult to enforce. This conflict, alongside biased per gene taxon sampling, is probably responsible for our difficulty in recovering some documented patterns of compositional evolution within angiosperms, such as increases in GC content in Poaceae (Serres-Giardi et al., 2012). Alternatively, the loci which most strongly express this and analogous patterns may not have been sampled in this dataset.

We highlight this problem not to single out these data or the original analyses as we recognize that many large-scale datasets inevitably face challenges when cleaning data. Instead, we want to underscore the importance of homology and orthology analyses in the construction of single gene alignments and gene trees. While errors like this may not greatly impact species-tree analyses, especially if they are mostly random between gene trees, they can dramatically limit the utility of these data for other analyses.

## Acknowledgements

We would like to acknowledge the importance of several discussions with colleagues including James Pease, Greg Stull, Jeremy Beaulieu and the Smith lab group. SAS was supported by a MICDE discovery grant and NSF 1938969 and 1917146. NWH was supported by the Woolf Fisher Trust.

## Author Contribution

SAS, CPF, and NWH contributed to the conception, programming, and writing of the manuscript.

## Data Availability

The alignments for both DNA and amino acid datasets are available through the resources of the original data release paper. The gene trees for DNA were generated as part of this study and are available from DataDryad. The code is available through github and sourcehut linked above.

