## Supplemental Tables and Figures for "Compositional shifts associated with major evolutionary transitions in plants"

### Supplementary Figure

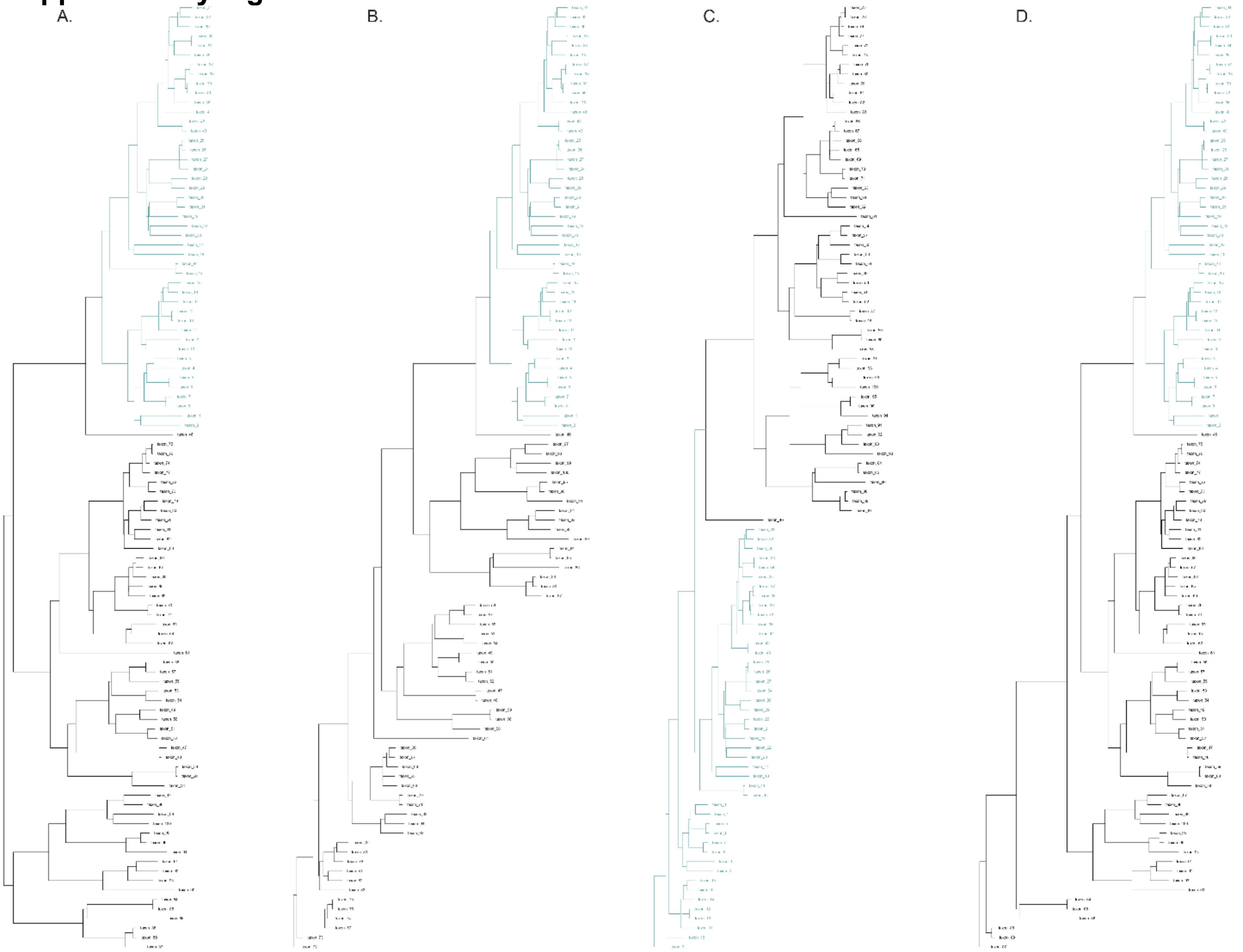


Supp. Figure 1. Given to complexities in the dataset (e.g., the sampling between gene regions, issues with monophyly), we were required to root in a greedy way (described in the Methods). The example here demonstrates that this is not a problem for our analyses. In this case, the blue clade has been simulated with differences in composition and is detected as divergent in each case. A) is the original rooting, B) is rooted as a grade, C) is rooted inside the shifted clade, and D) is an alternative rooting as a grade.

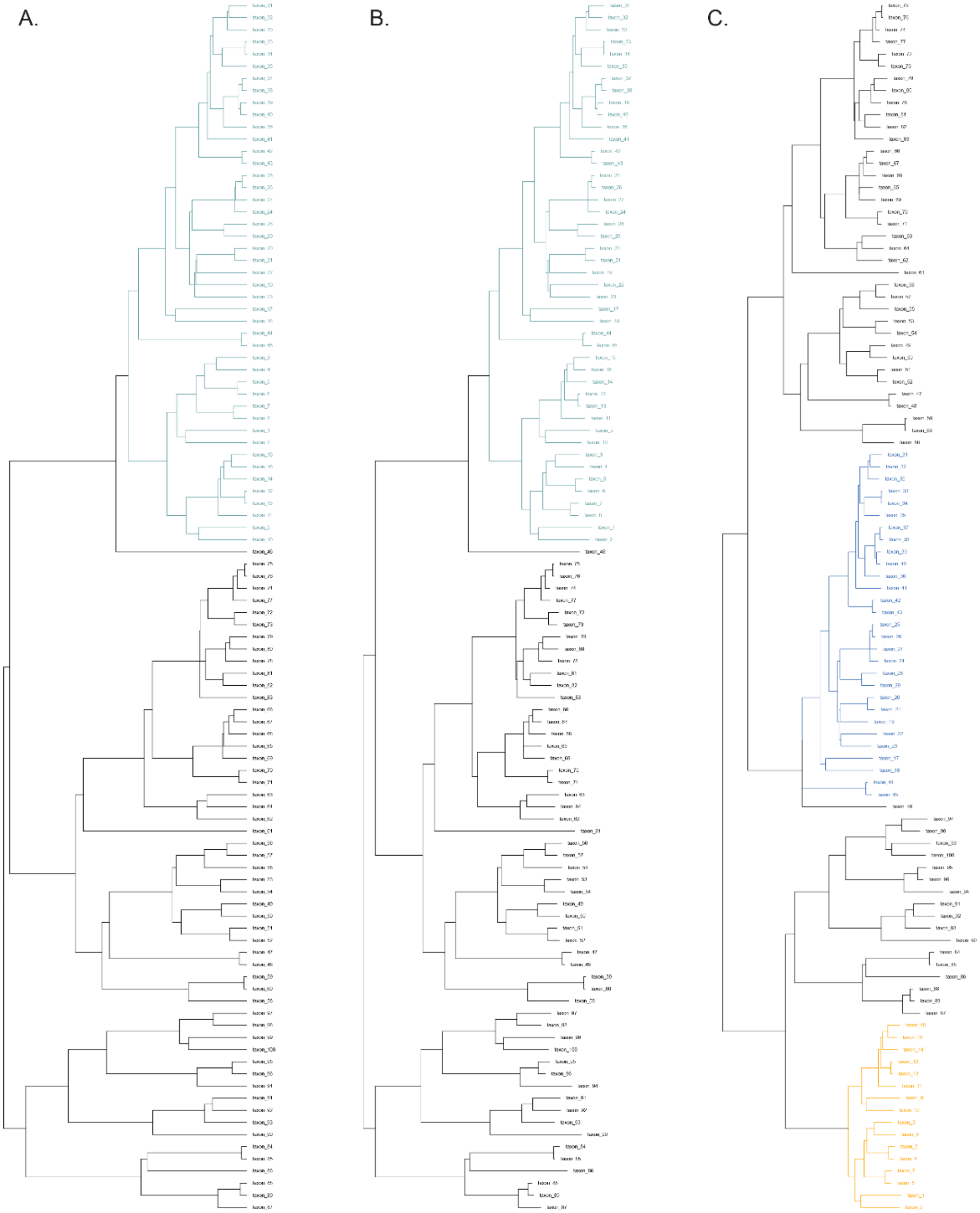


Supp. Figure 2. Compositional bias can potentially cause phylogenetic error. Phylogenetic accuracy is not the focus of this study and phylogenetic error can be caused by many sources. To demonstrate that the method of identifying shifts in the model is robust to phylogenetic error, we show a simulation of a rate shift reconstructed with the tree used for simulation (light blue in A), the reconstructed tree (light blue in B), and an incorrect tree (C). In C, the shifted clade falls in two parts of the tree. Nevertheless, the shift is detected in each subclade (shown as two colors).

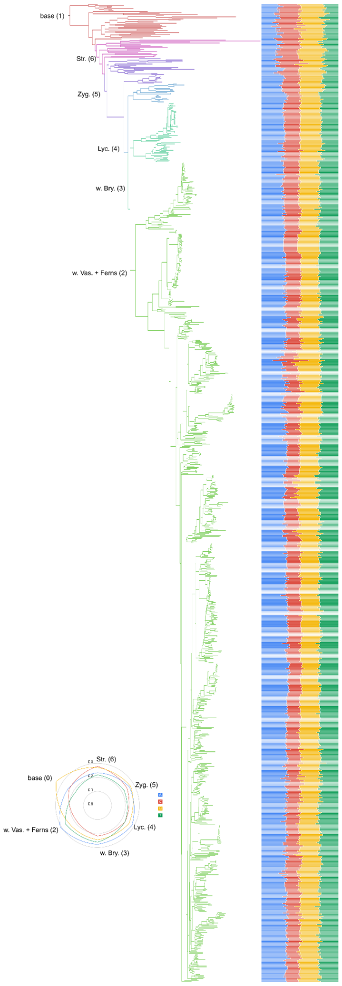


Supp. Figure 3. Composition of each tip for ortholog 5916 in the DNA dataset. The shifts are denoted on the tree with the values noted in the radar plot.

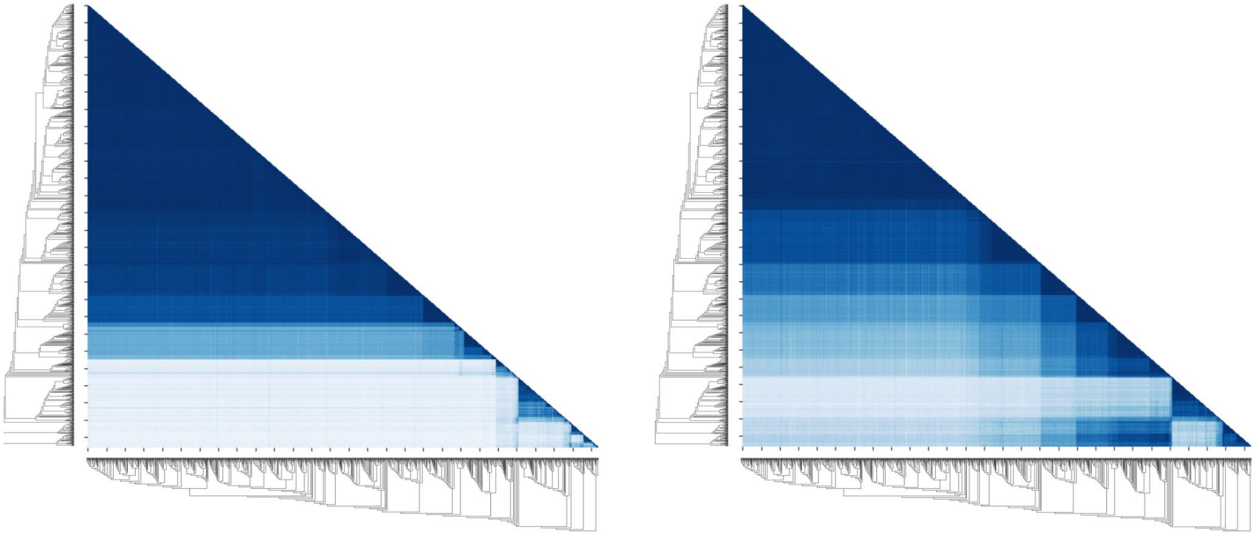

Supp. Figure 4. Pairwise heatmaps (left DNA and right AA) of shared models on the species tree (1kp citation).

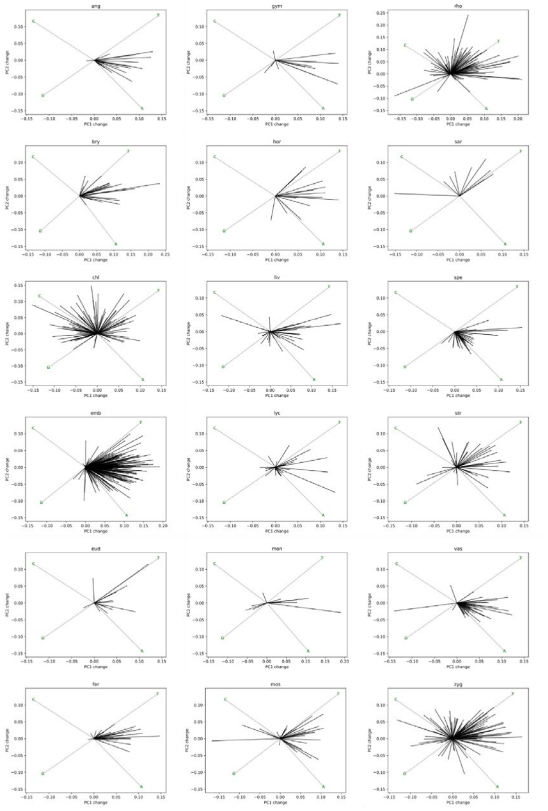

Supp. Figure 5. Vectors of composition shifts for DNA for major clades.

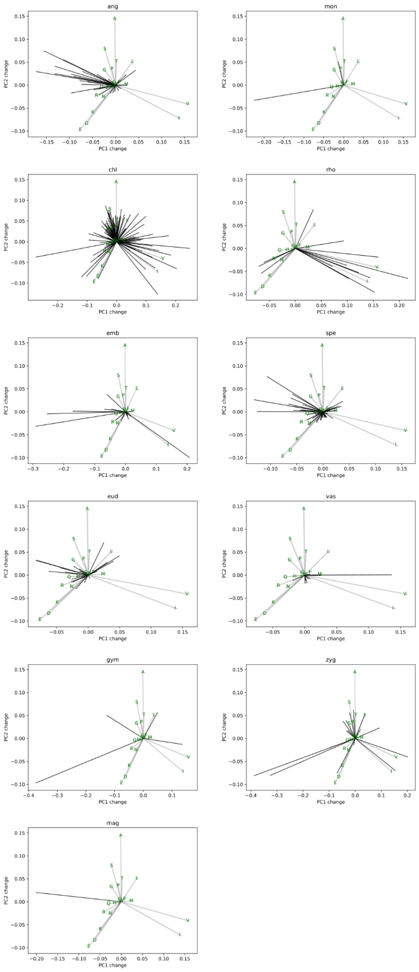

Supp. Figure 6. Vectors of composition shifts for AA for major clades.

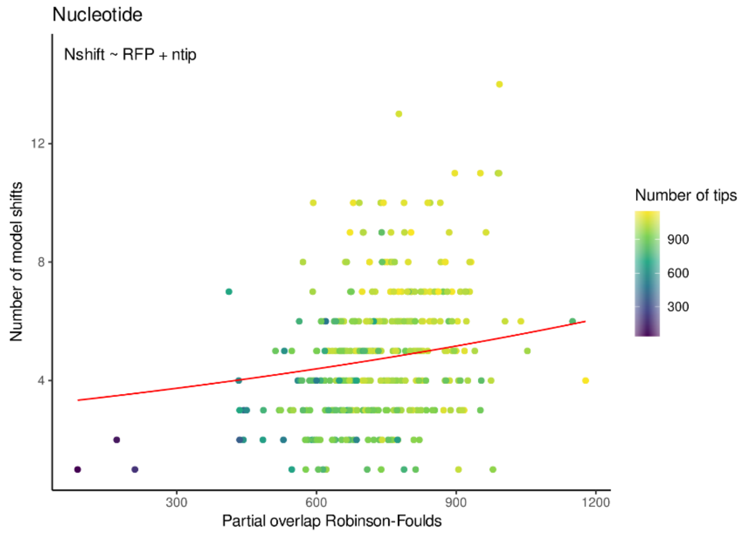


Supp. Figure 7. RF distances accounting for missing taxa calculated from individual DNA gene trees to the species tree plotted against the number of model shifts. Colors correspond to the number of tips in the tree. The line represents the the predicted number of shifts, holding number of tips constant (fitted with Poisson GLM in R). There is a weak positive relationship between increasing RF and number of inferred shifts, after accounting for tree size. There is also a relationship of increasing RF distance and size of the tree.

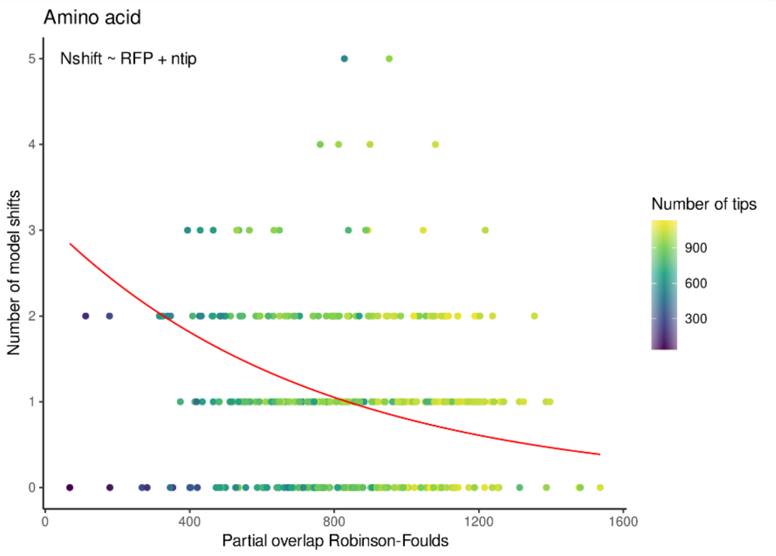


Supp. Figure 8. RF distances accounting for missing taxa calculated from individual AA gene trees to the species tree plotted against the number of model shifts. Colors correspond to the number of tips. The line represents the predicted number of shifts, holding number of tips constant (fitted with Poisson GLM in R). There is a negative relationship of increasing RF to inferred number of shifts. There is also relationship of increasing RF distance and size of the tree.
